# Disentanglement of Entropy and Coevolution using Spectral Regularization

**DOI:** 10.1101/2022.03.04.483009

**Authors:** Haobo Wang, Shihao Feng, Sirui Liu, Sergey Ovchinnikov

**Affiliations:** FAS, Division of Science; JHDSF Program, Harvard University, Cambridge, MA,USA,02138; Institute of Image Processing and Pattern Recognition, Shanghai Jiao Tong University, Shanghai, China 200240

**Keywords:** coevolution, entropy, spectral regularizer

## Abstract

The rise in the number of protein sequences in the post-genomic era has led to a major breakthrough in fitting generative sequence models for contact prediction, protein design, alignment, and homology search. Despite this success, the interpretability of the modeled pairwise parameters continues to be limited due to the entanglement of coevolution, phylogeny, and entropy. For contact prediction, post-correction methods have been developed to remove the contribution of entropy from the predicted contact maps. However, all remaining applications that rely on the raw parameters, lack a direct method to correct for entropy. In this paper, we investigate the origins of the entropy signal and propose a new spectral regularizer to down weight it during model fitting. We find the added regularizer to GREMLIN, a Markov Random Field or Potts model, allows for the inference of a sparse contact map without loss in precision, meanwhile improving interpretability, and resolving overfitting issues important for sequence evaluation and design.

## Introduction

Billions of years of evolution of natural selection have produced an astronomical number of diverse protein sequences. By comparing the sequences to each other, it has become possible to model the evolutionary constraints important for protein structure and function. Since it was shown the covariance patterns observed in a multiple sequence alignment (MSA) of homologous proteins are related to structure (1), models have been developed to automate the extraction of this coevolutionary signal for protein structure prediction and design.

The latest class of models, designed to disentangle direct from indirect coevolution (2) include Markov Random Fields (MRF) or Potts Models. The parameters of these model have been inferred using a plethora of methods such as GREMLIN (3), plmDCA (4), bmDCA (5), PSICOV (6) and mfDCA (7). This also includes the most recent low-rank reparametizations such as restricted Boltzmann machines or variational autoencoder(8), and self-attention-based models that share MRF parameters across protein families(9). The parameters from these models are used for protein structure prediction (10–14), protein-protein interaction prediction (15–17), protein design(18–20), mutation effect prediction(21, 22), and protein sequences alignment and homology search (23–26). The result of these models is typically two sets of parameters. One is an L x K matrix modeling the conservation and entropy, where L is the length of the protein sequence and K is the number of amino acids plus gap. The other is an L x K x L x K tensor modeling the coevolution. For contact prediction, the 4-dimensional coevolution tensor is reduced to an L x L matrix by taking the norm of each K x K matrices (Figure 1), followed by a low-rank correction procedure. The matrix represents the strength of the residue-residue interactions within the protein. The low-rank signal was shown to be highly correlated with entropy (27), indicating an entanglement of entropy and coevolution signal. Methods to correct for this include APC (28), LRS (29) and BND (30). Among them, Average Product Correction (APC) is widely used to boost contact accuracy in almost all of the coevolution models. For instance, in bmDCA, a model designed to recapitulate all the pairwise frequencies observed in the natural MSA, and most recently transformer-based models designed to share parameters across models, still rely on APC to get better contact prediction (9, 31). Though the models and loss functions are becoming more complex, they are still unable to disentangle the signal in the raw parameters of the model.

**Fig. 1.**
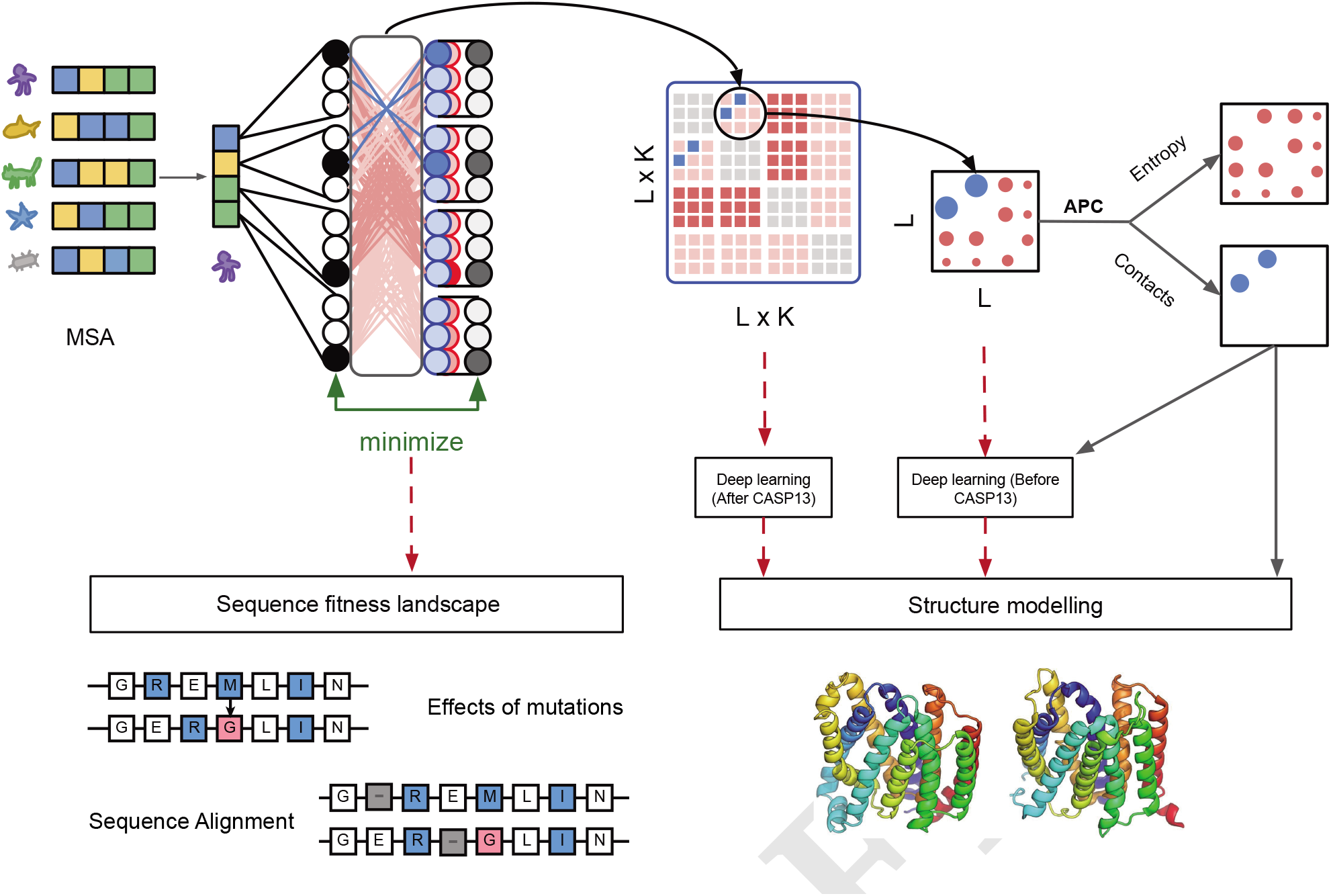
Introduction of entropy issue. MSA is fed into MRF model to reconstruct the input sequences back. The model contains a tensor modeling the coevolution (L x K x L x K), and a matrix modeling the conservation and entropy (L x K). Usually, the 4 dimensional tensor is reduced into an L x L matrix, then APC is applied to separate contacts from entropy signal. The contacts can be further used as constraints to model the structure. Prior to CASP13, the full 4D tensor was not widely used as input to deep learning, instead a reduced L X L matrix was used. The coevolution model can also be applied to predict the effects of mutation and align sequences. But it is worth noting that entropy signal (indicated in red) is only separated by APC method at the reduced matrix level. At the 4D tensor level, the coevolution signal and entropy are still entangled. The entropy issue might hurt the performance of all related applications through the dashed red lines.

For applications besides contact prediction, the full coevolution tensor, without entropy correction, is used as input to deep learning methods like direct structure prediction protocol Alphafold(14), protein design (19, 20), mutant ranking (21, 22) sequence alignment and remote homology detection (23–26) (Figure 1). Given the current correction methods, such as APC, only work for the L x L matrix, it remains unclear whether the entropic “noise” also affects these applications or not. Similar to how the entropy correction on the L x L matrix improves the interpretability and the accuracy of contact prediction, we reason a correction or regularization at the coevolution tensor level should also improve it’s down-stream application. Understanding the biophysical meaning of APC and how to apply the correction within the model rather than post correction remains a fundamental question in the coevolution field.

In this paper, in addition to confirming the low-rank signal removed by APC is correlated with residue’s information entropy (27, 28), we find that it also correlates with the dominant eigenvector of the 2D contact matrix(32). Based on these observations, we modify GREMLIN’s pseudolikelihood objective to include a spectral regularizer over the pairwise coevolution parameters. The parameters of this model are tested on both the task of contact prediction and sequence design tasks. For unsupervised contact prediction, using the norm of the coevolution tensor, the result shows that it can achieve almost the same accuracy compared with APC. For supervised contact prediction, contact accuracy improves when the coevolution tensor is used as input to a logistic regression model. For sequence design, we find the regularization improves model interpretability, revealing more biophysical details of each amino acid pair, potentially allowing for rational sequence design. Furthermore, we find the resulting Hamiltonian(or inferred energy) better correlates with protein stability.

## Results

### Recap: Markov Random Field

The characters of each string in the multiple sequence alignment are one-hot encoded. The data matrix **X** ∈ ℝ^*N* × *L* × *K*^ has *N* sequences, each sequence is of length *L* and each position in the sequence can have *K* different types of amino acids. In MRF model, one body term **B** ∈ ℝ^*L*×*K*^ and two body term **W** ∈ ℝ^*L*×*K*×*L*×*K*^ are used and the Hamiltonian term is written as 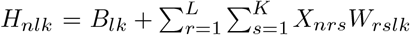 And pseudolikelihood method is used to approximate the partition function.

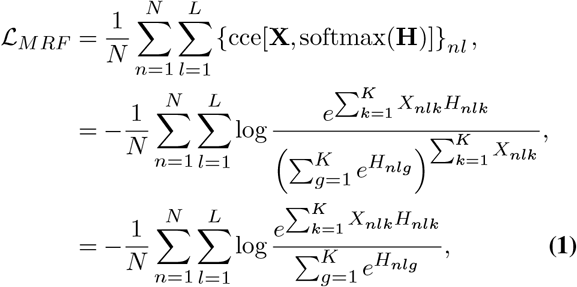

### APC is equivalent to removing the first eigen-mode of the L x L matrix

The contact matrix **M** ∈ ℝ^*L*×*L*^ is derived from the Frobenious norm of **W**.

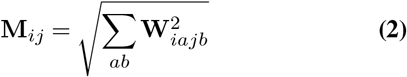

We use **p** to denote the sum of **M** per column/row, that is, **p** := ∑ _*j*_ *M*_*ij*_ = 1M ∈ ℝ^1 × *L*^. The background signal can be written as **p**^*T*^ **p**/∑_*i*_ *pi* (noted as AP term). The corrected matrix is denoted as **C**, thus we have,

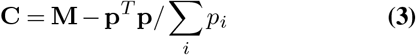

The Perron–Frobenius theorem (33, 34) says that for a non-negative real square matrix, there exists a non-negative dominant eigenvalue. This means for the matrix **M** there exists a dominant eigenvalue that can be approximated by power iteration method. We demonstrate that APC is approximating the first eigen-component by initializing a **1** vector (mathematical proofs can be found in supplementary information). Thus, the APC can be also written as

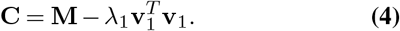

where *λ*_1_, **v**_1_ are the dominant eigen value and eigen vector of **M**.

### APC signal and model overfitting

#### Spectral Regularization

To introduce the average product correction (APC) during training process, we propose to remove the first eigen-mode of the **M** at gradient level. This indicates the regularizer can be written as the integral of the first eigen mode. So during training, the first eigen-mode will be down-weighted in each step.

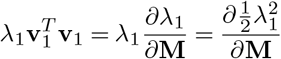

Combined with hyper-parameter, we define the final regularizer LH as 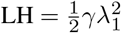, where *γ* is the hyperparameter. We can approximate *λ*_1_ by APC (details in supplementary information).

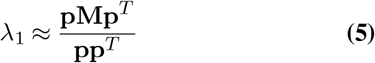

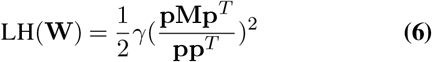

A multiple sequence alignment (MSA) is a collection of evolutionary related sequences. The relationship between positions (or columns) of the MSA are due to structure constraints and relationship between sequences (rows) are the phylogenetic signal. These two signals can be entangled (35, 36), especially when the sample size is low. Positions with high entropy just by change may appear to be “coevolving”. This is evident by looking at the normalized **p** vector from the coevolution matrix **M** (averaged column/row). The **p** vector has been reported to be linearly correlate with the square root of entropy (27). Even with this observation it remains unclear how to disentangle them within the model.

An obvious solution is to use a sparse regularizer, such as Block L1 (LB) (3), but this was surprisingly found to result in less accurate contact prediction compared to L2 with APC. To better understand this phenomenon, we reexamine the correlation between **p** and entropy with respect to the depth of MSA for a protein family (PDB: 3CNB, chain: A) with at least 20K sequences. With few sequences in MSA, the correlation between **p** and square root of entropy is almost linear. When the number of sequences increases, the Pearson correlation between these two vectors starts decreasing (Figure 2A). To evaluate the over-fitting issue, we randomly split the MSA into training sets and test tests. Then, we checked the distribution of loss in these two sets, and quantify the over-fitting in MRF model using the Kullback–Leibler divergence (Figure 2B). The KL divergence shows that the model tends to be overfitting when it does not have enough sequences, consistent with previous reports(37). We also see the fraction of variance explained by the first or dominant eigen-mode of **M** to decrease with more sequences. Yet even at 20 thousands sequences, 90 percent of **M**, (Figure 2C) is dominated by the largest eigen-mode, which average product correction (APC) approximates via first power iteration (see details in supplementary material). The sparse structure information only explains less than 10 percent of **M**. Based on this observation we reasoned suppressing the first eigen-model in **M** can be a prior to strengthen the sparse structure information.

**Fig. 2.**
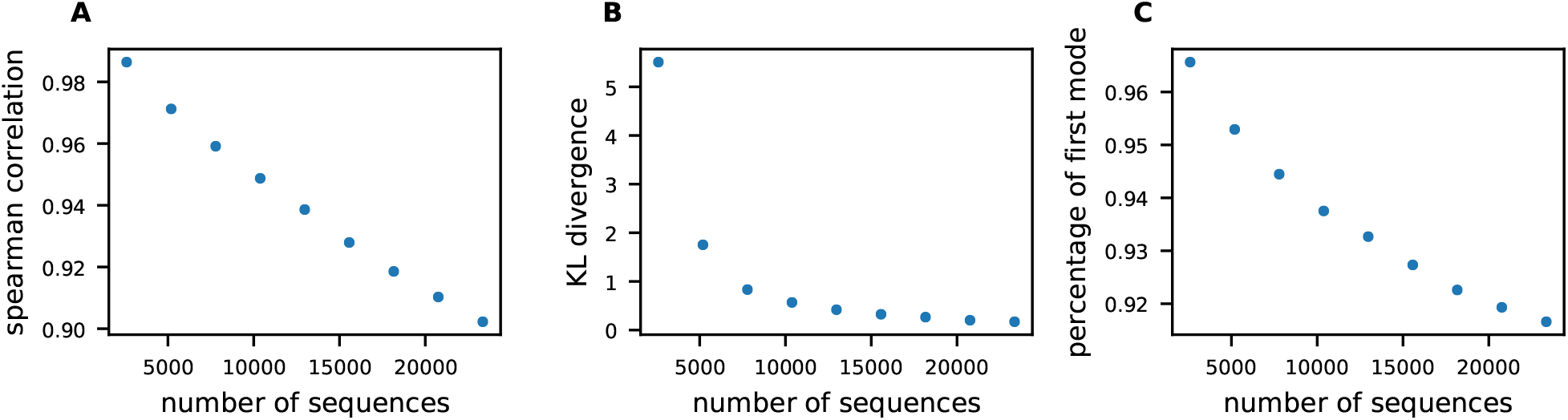
APC signal and model overfitting. Figure A,B and C, the x axis is number of sequences. The y axis in Figure 2A refers to the Pearson correlation between square root of entropy and **p** vector. The y axis in Figure 2B refers to the KL divergence between the loss distribution of training set and test set. The y axis in Figure 2C refers to the percentage of first eigen mode in **M** matrix.

Inspired by APC, we propose a spectral regularizer, named as LH (abbreviation of Henri Lebesgue) regularizer, designed to suppress the first eigen-mode in the pairwise parameter of the MRF during training at the gradient level. For visualization, as is shown in Figure 3, we analyzed the gradient of three different regularizers, L2, LH and LB. Instead of showing gradient of **W** matrix, we show the gradient based on **M** matrix. It is obvious that L2 method is re-scaling the matrix and has no contribution on removing the entropy or promoting the sparsity. LB has a constant gradient to all the elements. For LH, the gradient captures the **P** term, which means the entropy in the two body term can be effectively suppressed during training process.

**Fig. 3.**
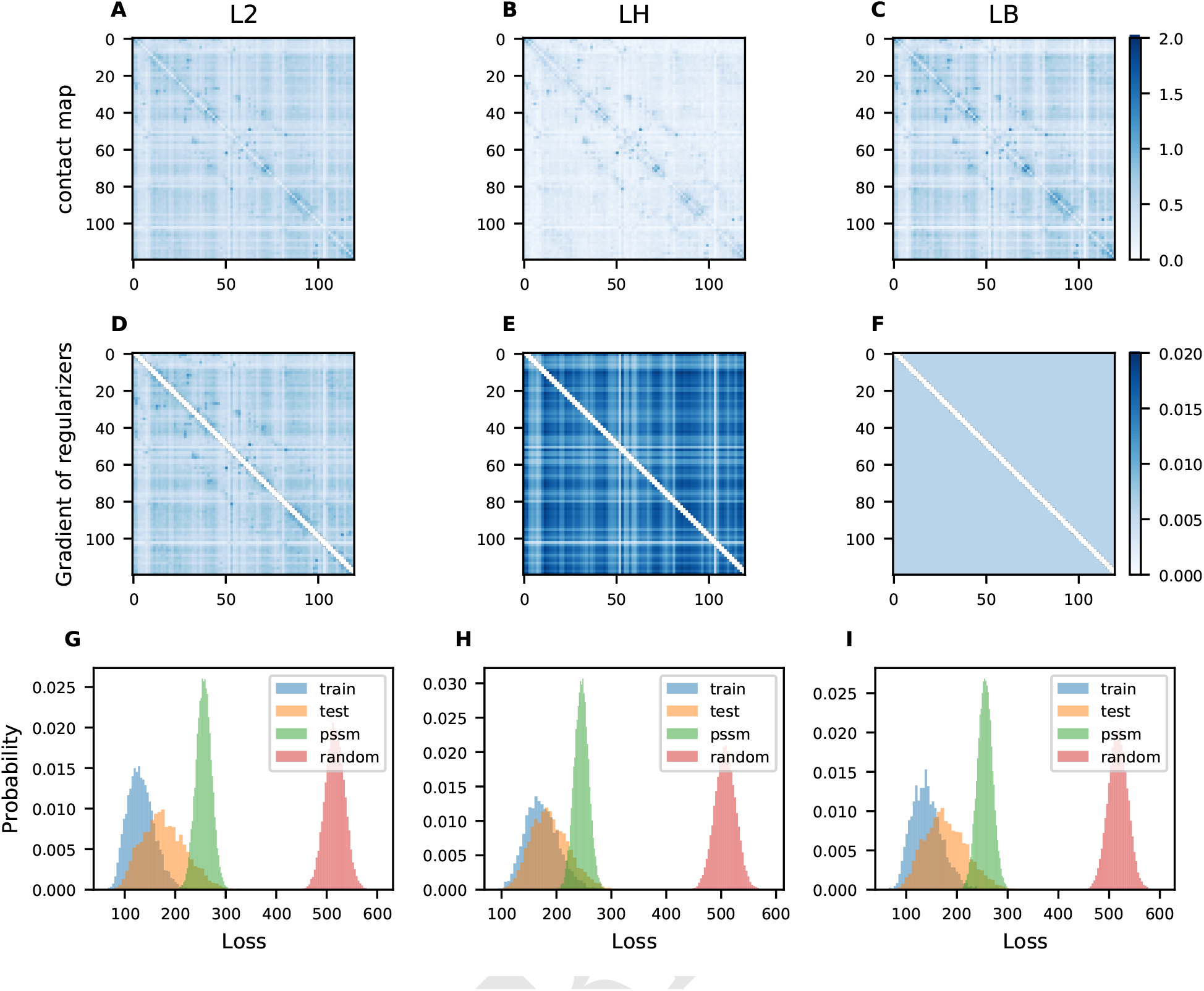
The effects of regularization type (L2, LH, and LB) on the sparsity of contact maps, gradient and overfitting. The first row (A,B,C) shows the **M**, with L2, LH, and LB regularization respectively. The second row (D,E,F) shows the gradient of these three regularizers. The third row (G,H,I) shows the distributions of reconstruction losses for the natural MSA training set (blue), test set (yellow), MSA sampled from a PSSM of the natural MSA(green), and MSA sampled from a random distribution MSA(red). Even though the loss distribution of the training set is well separated from the PSSM and randomly sampled sequences, for the L2 and LB regularized models the test set distribution does not overlap with the training set loss, indicating an overfitting issue. For the LH regularized model, the training and test set loss distributions have a good overlap.

As is shown in Figure 3, unlike the L2 or LB regularized parameters, the LH regularized parameters no longer exhibit the low-rank signal (vertical and horizontal lines) that require APC for the unsupervised contact prediction task, nor do the parameters show signs of over-fitting (differences in the loss or the negative pseudo-likelihood distribution of the training and test set). The same is observed across 9 more protein families with various depth (Figure SI1), LH regularized parameters consistently avoid the over-fitting issue.

To test the robustness of the loss or the statistical energy of a sequence (Hamiltonian), we first compute the reference Hamiltonian using a protein family (PDB: 3CNB, chain: A) with more than 20K sequences, using the parameters from three different regularization schemes. Next, we sub-sample the MSA to different depths, refit the parameters and compute the Spearman correlation to the recompute Hamiltonians. As seen in Figure S3 and Figure 4, the correlation between Hamiltonian with ref. Hamiltonian is more robust under the regularization of LH. As for L2 and LB, the correlation drops rapidly with the decrease of sequences, indicating a more severe over-fitting issue with less data.

**Fig. 4.**
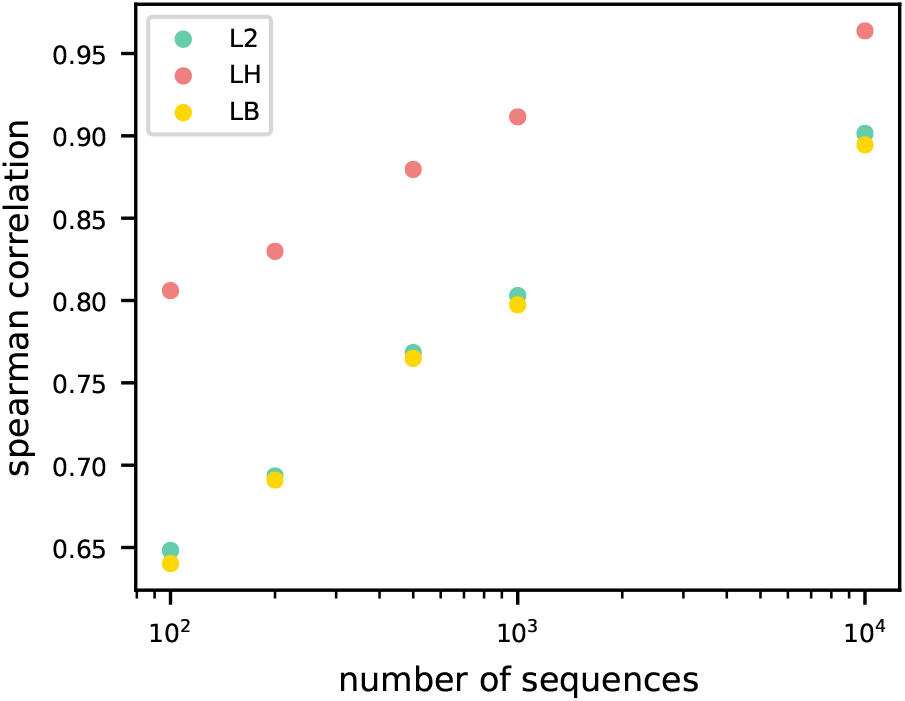
Robust recovery of the Hamiltonian with fewer sequences under LH regularization. The x-axis shows the number of sequences, the y-axis shows the Spearman correlation of the Hamiltonians from the original MSAs and a subset of MSAs with various number of sequences. All three methods are shown on this plot.

Furthermore, to demonstrate the disentanglement of coevolution and entropy (as measured by conservation), we use the parameters of the L2 and LH models to sample new sequences. Specifically, we sample sequences based on one-body parameters, two-body parameters, or the combination of both. If disentangled properly, the sampling procedure should require both terms for the PSSM (Position-Specific Scoring Matrix) of the sampled sequences to match the PSSM of the natural sequences. To test this, we sample sequences for a set of 553 protein using CCMGEN (27) (see methods). As shown in Figure 5, when using both one-body and two-body parameters (wb), the PSSMs of both LH and LB regularized models match. When using just the two-body parameter (w) for sampling, the PSSMs match for L2 but not for LH. The opposite is observed when using just the one-body term (b) for sampling, the PSSMs do not match for L2, indicating the entanglement of the entropy and coevolution signal in the two-body parameters, with the one-body term playing little role. For LH regularized models, the best correlation is achieved when both the one and two-body parameters are used, indicating the disentanglement of coevolution and entropy.

**Fig. 5.**
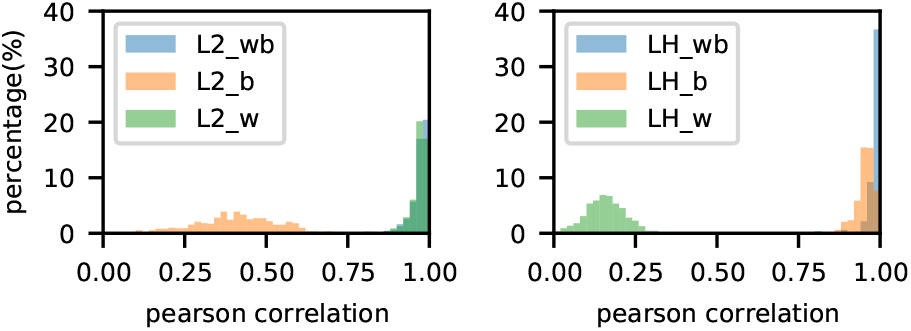
LH regularized parameters disentangle conservation and entropy from coevolution while L2 regularized parameters do not. The figure shows the distribution of Pearson correlation values across different proteins, comparing the PSSMs of the sampled sequences vs. the natural sequences. The sampling was done using just one-body (b), just two-body (w) or the combination of both one and two-body (wb).

Guided by these results, we tested this method with more protein related applications, such as contact prediction and sequence design, to see if disentangling entropy inherently helps enhance their performances.

#### Unsupervised and Supervised Contact prediction

For the contact prediction task, the contact precision is evaluated on a dataset of 383 proteins (see methods). Two separate matrices are evaluated, the Frobenious norm of **W**, denoted as ‘raw’ matrix and the average product corrected matrix denoted as ‘APC’ following Equation (3). As is shown in Figure 6A, with increased weight of LH regularization the performance of the ‘raw’ and ‘APC’ matrix become similar in contact precision and approach the precision of L2 regularized ‘APC’ matrix. Interestingly, with lower regularization weight, the ‘APC’ matrix of LH regularized model outperforms the original markov random field (MRF) model with L2 regularization (Figure 6C). For comparison, we also try Block L1 (LB), with increased weight the ‘raw’ performance improves, but does not reach the performance of L2 plus ‘APC’. Though both LH and LB regularizers improve the performance over L2, when comparing the ‘raw’ matrices (Figure 6B and Figure 6D), only the LH ‘raw’ approaches that of the L2 ‘APC’. In conclusion, the LH regularizer removes the need to do APC, while preserving the contact precision of the MRF model.

**Fig. 6.**
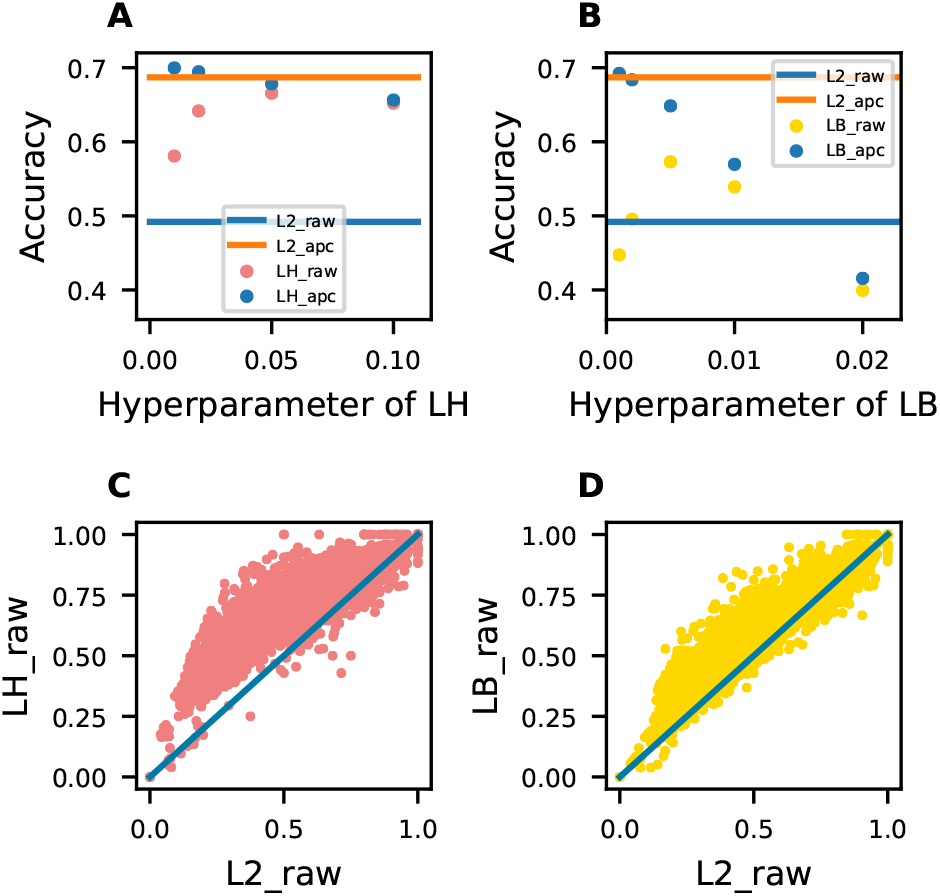
For unsupervised protein contact prediction, average product correction (APC) is no longer required under the LH regularizer. (A) The accuracy of L2 based method is set as baseline with two solid lines. Blue one is the performance without APC, and the orange one is the accuracy with APC. The x axis is the hyperparameter for LH and the red dots showed the raw accuracy of LH and blue dots with APC correction. (B) The accuracy of Block L1 is showed with different hyperparameter scanning. (C) The best raw performance from LH compared with L2 based method, Each point corresponds to a protein MSA, the axes indicate the accuracy of each method, defined as the average precision of the top L ranked contacts, L is the length of the protein. D) Performance comparison between L2 and LB.

Since all residue-residue interactions within a protein are governed by the same basic physical potentials we expect interacting residues pairs to share a limited number of K x K matrices. To test if LH regularized parameters better recover these, we apply principle component analysis (PCA) to the **W** matrix. PCA is a decomposition method, allowing distillation of important common or shared features. We transpose the **W** matrix to treat the K x K dimension as features and L x L as number of samples. The K x K dimension carries the biophysical meaning of amino acid interactions, and are expected to be revealed by principle components. We can see from the Figure S2, the LH regularizer requires fewer number of principle components to explain the same amount of data compared to L2 and LB. Current machine learning model usually use the **W** matrix as an input to predict the protein contact map or distance matrix for future structure prediction (12–14). We reasoned the distilled features may be more useful as inputs to machine learning input and can be trained as input of a supervised machine learning method. To test this hypothesis we trained a simple logistic regression neural network (denoted as “NN“) similar to the one in (9) to learn a weighted sum of the K x K matrix for contact prediction.

We evaluate the performance of the 9 methods and show the precision on the top L contacts in Figure 7. For contact definition, we use ConFind (38) with contact degree cutoff of 0.01 and sequence separation of >= 6 Angstrom. The 9 methods include the pairwise combination of regularizers (L2, LH and LB) and the three post-processing methods (raw, APC and NN). From the results, we can see that LH-based inputs are more robust and have higher precision, suggesting the utility of the denoised or distilled features. Given these results, we expect the features extracted in the context of LH regularization will also improve the results in more advanced deep neural networks.

**Fig. 7.**
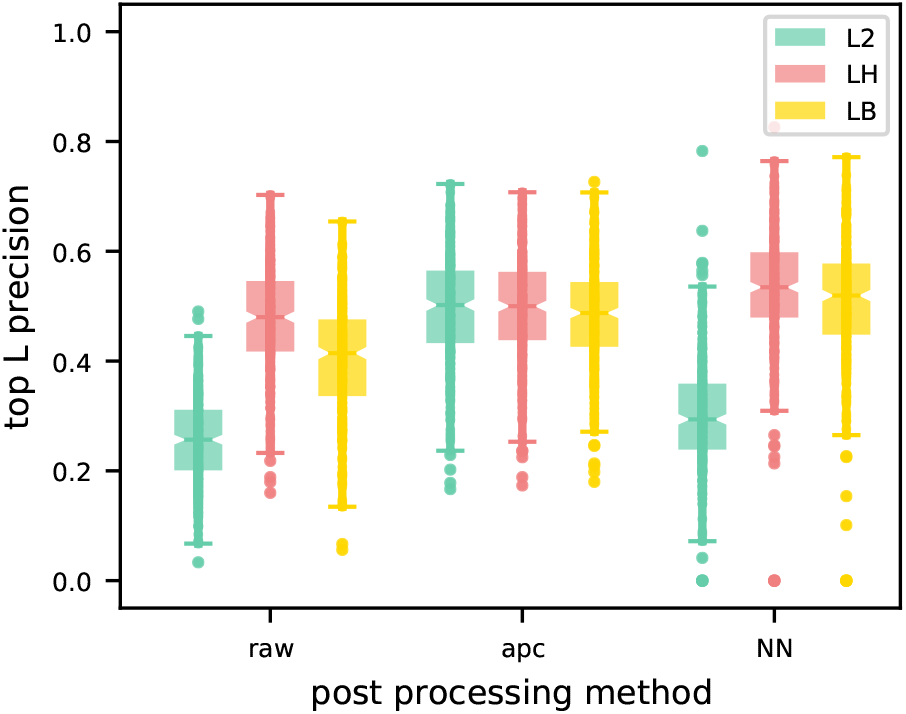
For protein contact prediction, three different way of contact extraction are applied to all three methods,L2(green), LH(red) and LB(yellow). Besides the raw and APC method, we treated the **W** matrices as inputs of a simple regression model. The top L precision is used to curve the performance, L is the length of the protein.

#### Sequence design

Evolution-inspired sequence design is already implemented in some generative models such as Markov Random Field model(19), bmDCA(20), or energy mixture model(39). These models allow the calculation of the Hamiltonian as a statistical energy (as described in methods, see Markov Random Field) describing the protein thermal stability or fitness landscape. Using Monte Carlo to search sequence space and get the lower Hamiltonian is a current strategy to design sequences. Unfortunately, in these models, coevolution, entropy and phylogeny signals are entangled under the L2 regularizer. Since protein stability is thought to be due to structural constraints, we hypothesize a sparse model that only models the coevolution signal maybe be more prepredictive of protein stability.

To test if sparse models are more predictive of stability, we explored the stability of a series of reported designed sequence variants with labeled experimental data. We evaluated the performance using the Spearman’s rank correlation coefficient *ρ* between Hamiltonian and the melting temperature (denoted as ‘Tm’). As shown in Figure 8, we can see for the GA and GB binding domains of streptococcal protein G, the Hamiltonian from the sparse LH model is well correlated with the Tm. In Protein GA, the LH achieved performance with 0.76, slightly better than L2 model of 0.73. While in Protein GB, the LH have stable performance of 0.86, but the L2 drops down to 0.47. These two analysis demonstrated that LH based method might help design more stable proteins.

**Fig. 8.**
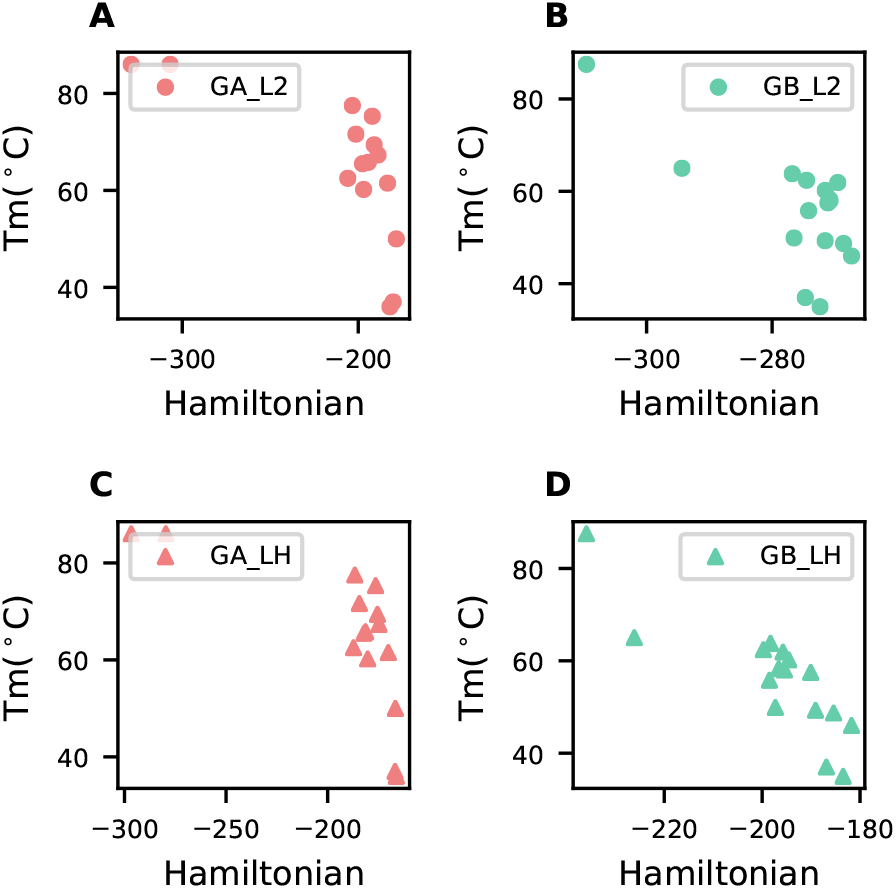
Reanalysis of Spearman correlation between Hamiltonian of GA/GB and published folding temperature(19). The L2 and LH regularizer are applied to curve the Hamiltonian. The x axis is the Hamiltonian and the y axis is the reported Tm. each point is a designed protein sequence.

#### The effects of spectral regularization on phylogenetics

Beyond the entropy signal, there is also the phylogenetic signal which is thought to be entangled in the coevolution matrix. To test this, we use the chorismate mutases dataset from (20) to fit three generative models and sample new sequences. Unless the sequences are explicitly sampled along a phylogeny (27), we would expect independently sampled sequences to be devoid of low-rank signal representing relationships or clusters of sequences. Sequences sampled from L2 regularized models bmDCA and GREMLIN fully and partially preserve, respectively, the low-rank structure when projected onto the two largest principle components of the natural MSA. For LH sampled sequences, the signal is gone, indicating that the phylogenetic bias is now suppressed as shown in Figure 9.

**Fig. 9.**
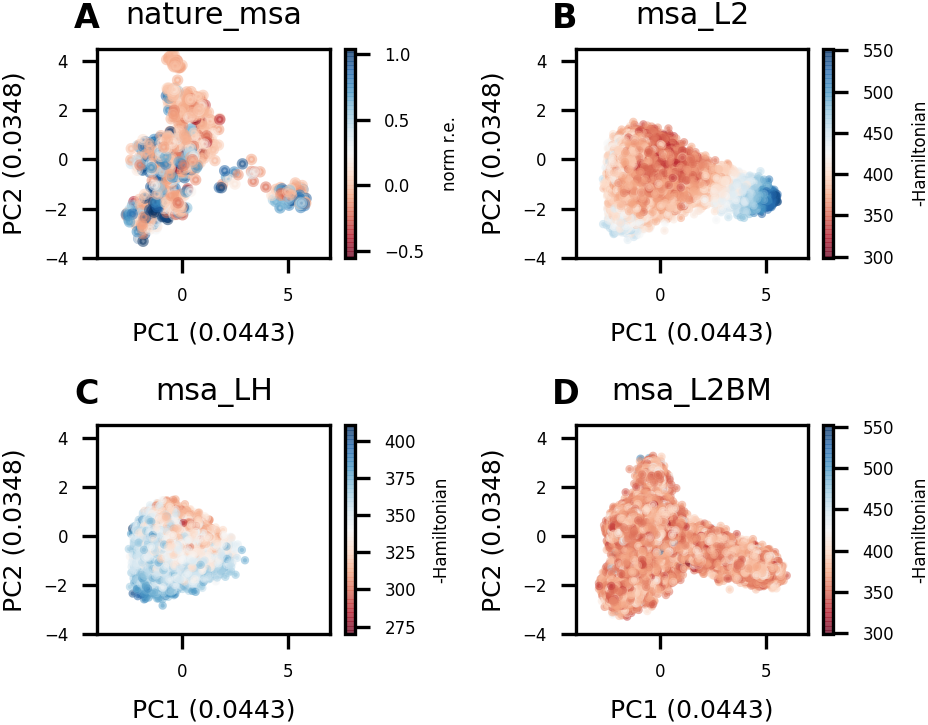
PCA analysis of MSAs to check about the sequence relations. all sequences are projected into PC1 and PC2, the first plot is colored by norm r.e.(fitness) and the rest plots are colored by negative Hamiltonian. Blue couples with positive fitness.

Though this is theoretically a good result, it can be problematic for sampling of functional sequences, if the low-rank signal represents functional clusters as expected for an MSA that is a mixture of paralogs and orthologs. The low-rank signal maybe useful for discriminating functional from non-functional sequences, if different clusters represent different function. To test that the LH regularized model is still able to discriminate between working and not working designs, we retrain the models using a subset of natural sequences experimentally determined to have the desired activity (denoted as “norm r.e.”). As shown in Figure S6, for all 3 models, we find the designed sequences from (20) can be easily separated into working and non-working by their computed Hamiltonian. To confirm that the sparse coevolution signal is present, we compute the Mutual Information (MI) contact maps of the L2, LH and bmDCA (denoted “L2BM“) sampled sequences. Analyzing the contact maps qualitatively, we see low-rank signal (vertical and horizontal lines) in the MI matrix for L2 and bmDCA sampled sequences, yet little signal in the LH sampled sequences. To confirm t he e ntropy s ignal i s preserved, we compared the marginal entropy of the sampled sequences to those of the natural MSA and see a strong correlation (Pearson r=0.95), suggesting the entropy is still captured by the LH regularized model but disentangled from the pairwise term.

These results suggest a path forward towards development of a unified model that accounts for both coevolution and phylogeny. LH maybe useful to disentangle the covariance due to phylogeny vs. coevolution in a model that parameterize these two signals.

#### Interpretability

LH improves interpretability of single sequence contacts in coevolutionary models. For a given sequence, we expect the parameters of MRF models to represent physical potentials of interacting residues, yet when we analyze the L2 or LB regularized parameters for a given sequence, we see vertical and horizontal lines of indirect correlations. For LH regularized parameters the signal is sparse (figure S 6), a nd r eveals a ttractive o r r epulsive interactions, unlike the L2norm used to represent the average signal across a protein family. As a case study, we analyze adenylate kinase (40). From the figure 1 0 w e c an s ee, A NC1 i s the most ancient sequence and has the melting temperature of 89 Celsius, with ANC4 only 75 Celsius. From the detailed single sequence contact analysis we see that ANC4 forms two strong negative or repulsive contacts compared to ANC1. The residue pairs between 23 and 209, changed to a repulsive lysine(K)-lysine(K) interaction in ANC4, while it is a stable lysine(K)-glutamic acid(E) salt bridge in ANC1.

For the residue pair 19 and 202, the salt bridge is also broken by mutating the aspartic acid(D) to asparagine(N). More details are shown in figure 10. From AN1 to *B. marinas*, besides breaking the 19-202 salt bridge, a *π*-cation interaction Phenylalanine(F)-K is broken by mutation of a neural interaction F-Glutamine(Q), which decreased the interactive energy, thus also weakened the stability of Adk from *B. marinas*. From ANC1 to *B. subtilis*, besides breaking the the cation interaction of R-Y to R-S, the well structured zinc binding sites Cysteine(C)CCC are mutated to CCCD, which not only shows a series of negative signal in metal binding region, but also might have an important effect to decrease the stability. This effect has not been reported in the published paper(40), and the zinc binding sites are reported to be highly correlated with stability recently(43). This qualitative analysis demonstrates the power of LH regularization in increasing the interpretability of coevolution of interacting residues and may help biologists to rationally analyze the effects of mutation for sequence design.

**Fig. 10.**
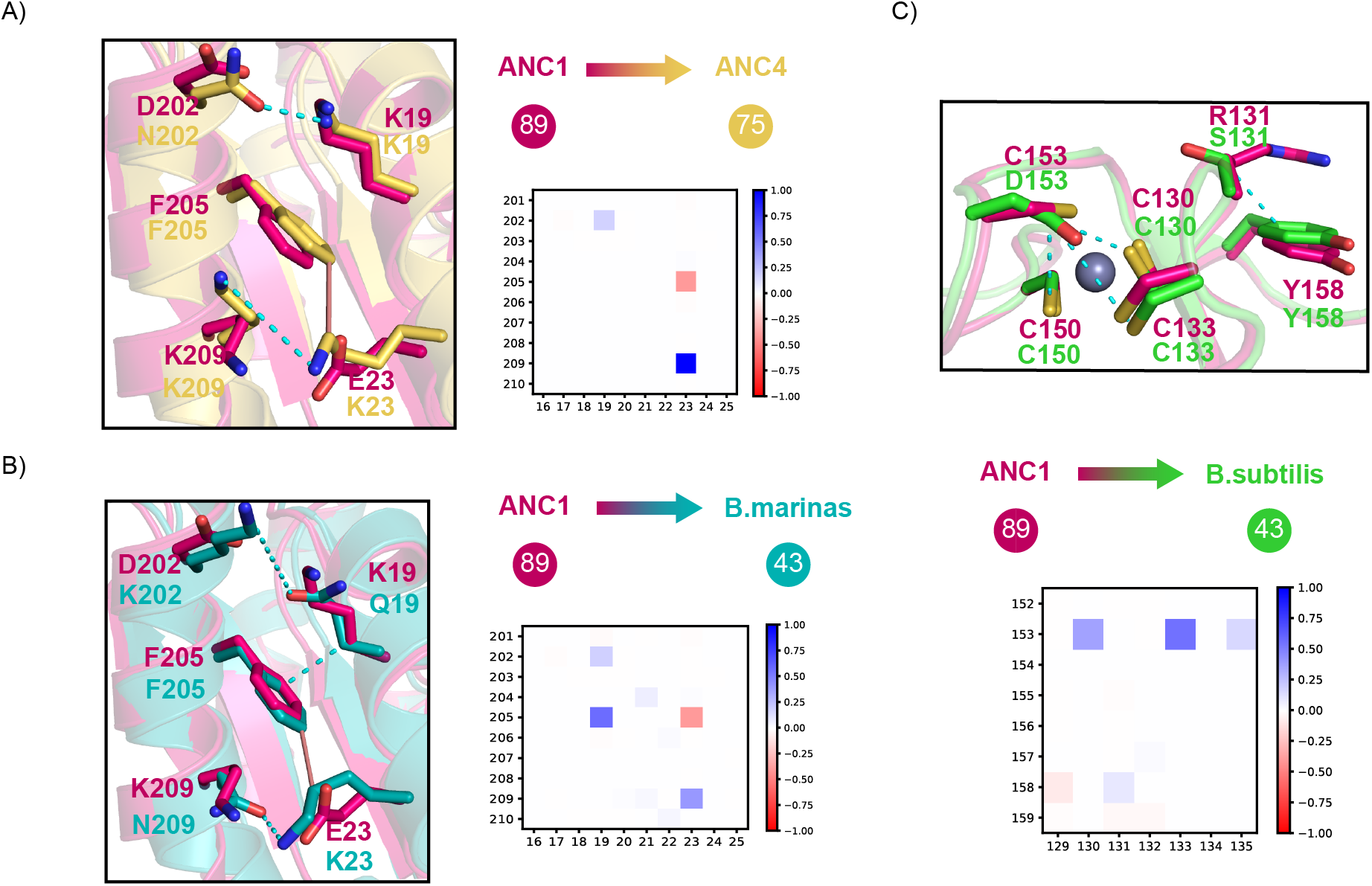
Structural reanalysis (40) of thermal stability from A) ANC1 to ANC4, ANC1 to *B. marinas*, ANC1 to *B. subtilis* using published crystal structures(40–42). The right panel showed the difference of coevolutionary coupling strength of two different proteins. The more unstable the right pairwise interaction is, more blue the pattern shows. The solid red lines represent high thermal stability, and the dashed dotted line weakened the interactions for the right protein.

## Discussion

Inspired by mathematical reinterpretation of average product correction (APC), we developed a spectral regularizer that penalizes the largest eigen-mode of the pairwise parameters of the markov random field (MRF) during training. This we find largely removes the need for low-rank post-correction. For the unsupervised contact prediction task, we find the extracted contact map no longer requires APC, largely matching the performance of L2 regularized methods with APC. In our LH model, the pseudo-likelihood (or self-supervised objective) is enforced by setting the diagonal part of **M** matrix to zero, this adds a constraint that the summation of eigenvalues equals summation of diagonals, which is zero. Suppressing the largest eigenvalues may also be distorting the rest of the eigenvalues. For APC, this removes the largest eigen-mode, and does not affect the rest eigen-modes. This may explain why some overfitting remains and the performance does not exceed that of L2 regularized models with APC. Further development maybe needed to limit the regularization to just the largest eigen-mode.

That being said, we find these parameters to be less over-fit in sequence reconstruction task, and thus may generalize into unknown space. The LH based **W** matrix is a better input for supervised learning, which might help to predict protein contacts in more complex supervised models. Further, we applied LH based MRF model to analyze the designed sequences from various examples, which all show strong correlations between experimental stability/fitness data and the Hamiltonian from our method. What’s more, it reveals more evolutionary patterns and increased the model interpretability. Taking all these into account, the LH based model is guided by structural principles and can further explore structural related applications. The removal of entropy and low-rank signal representing phylogeny from the two-body term opens the door to shared parameters and explicitly modeling phylogenetics. We suspect the two-body could be described by a limited number of 20 by 20 matrices representing biophysics, and these can be shared across protein families. Due to entanglement, we suspect prior work failed to improve contact prediction when sharing parameters across protein families (31) or explicitly modeling of phylogeny on real data (44, 45). With LH regularization, it may be worth revisiting these problem and approaches. Recent deep learning methods like VAEs (46, 47), BERT(48, 49), MSA transformer(50)), RoseTTAFold(51) and AlphaFold2(52) while do not explicitly parameterize MRFs, they are thought to learn them via the hidden parameters. These model optimize an approximation of the pseudo-likelihood function called self-supervision or masked-language-modelling (31). We suspect the Jacobian of these models can be computed (8) and regularized with LH to promote sparsity in the hidden representations.

## Materials and Methods

### Regularization

L2 and Block L1(LB) regularizers can be presented as follows,

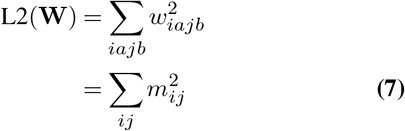

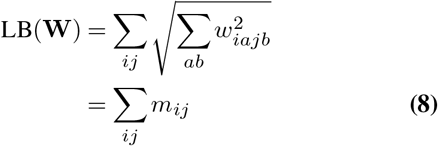

### Dataset and preprocessing

A dataset of proteins from the PDB database, along with their multiple sequence alignment (MSA) were collected from paper(53). To make the dataset consistent, a diverse subset of 383 proteins were selected that contained at least 1K sequences and sub-sampled to 1K sequences, as described in (54). For protein design, the dataset includes protein thermal stability data from GA and GB binding domains of streptococcal protein G (19) and the fitness measurements of chorismate mutases (20) and the Adenylate kinase sequences (40). The latter sequences were generated using an ancient sequence reconstruction approach.

For the disentanglement of entropy and coevolution experiment, 553 short proteins with at least 100 sequences in the MSA were collected from the PDB database. Considering the sampling efficiency of CCMGEN, the dataset was restricted to a maximum length of 80 amino acids. For the 553 proteins, we ran the TrRosetta’s HHblits protocol to generate the multiple sequence alignment. To summarize, the protocol starts with a search against the Uniclust30 sequence database. If fewer than 128 sequences are found at e-value of 1e-3, the MSA is further enriched using the BFD database. Once the parameters are fit using GREMLIN, CCMGEN (starting with a random sequence with burn-in of 1000) is used to sample 2000 new sequences. The datasets are deposited here (https://github.com/sokrypton/GREMLIN_LH).

### Supervised learning

To convert the two body term in MRF model into the 2 dimensional contact map, we make use of three conversion methods which are denoted as ‘raw’, ‘APC’, and ‘LR’. The ‘raw’ is **M** from Equation (2), and ‘APC’ is **C** from Equation (3). The ‘LR’ is the output following logistic regression fitting described in paper(9). We train the logistic regression on the curated dataset(54) using 5-fold crossvalidation, which contains 383 proteins. The dataset is first split into five equal parts. Five separate models were trained, for each model, 1/5th the data is selected as the test set and the remaining as the training set. In this way, the logistic regression model can be trained and evaluated on all the 383 proteins. The logistic regression model is L2 regularized with the coefficient of 1e-5. The learning rate is set to 5e-3 and the Adam optimizer(55) is employed to optimize the loss function.

The performance of 9 methods (pairwise combination of three regularization method: L2, LH and LB and three contact extract methods: ‘raw’, ‘APC’, and ‘LR’) are evaluated on the 383 proteins with contact precision of the top L predictions Figure 7.

## ACKNOWLEDGEMENTS

The authors thank Dr. Justas Dauparas and Dr. Young Lee for kindly and helpful discussion, thank Dr. Pengfei Tian, Dr. Christopher Wilson and Dr. Dorothee Kern for offering original data for further analysis. S.O. is supported by the John Harvard Distinguished Science Fellows Program within the FAS Division of Science of Harvard University. Research reported in this publication was supported by Office of the Director of the National Institutes of Health under award number DP5OD026389. The content is solely the responsibility of the authors and does not necessarily represent the official views of the National Institutes of Health.

## Proof

### APC approximately equals removing the largest eigen mode

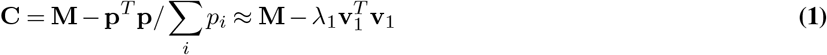

**M** is the Frobenious norm of the parameter inferred from Markov Random Field, **C** is the APCed matrix, **p** presents the sum of the column of the matrix **M**, *λ*_*i*_ and *v*_*i*_ is the i-th eigen component from the **M**.

Sum of column or row can be written as an operation with multiply **1** vector. We define **1** = [1, 1, …, 1] ∈ ℝ^1×*L*^, and there exists a set *α*_1_, *α*_2_, …*α*_*n*_ that can rewrite **1** = *α*_1_**v**_1_ + *α*_2_**v**_2_ + … + *α*_*n*_**v**_*n*_. The *α*_*i*_ is the **1** vector that projection to the eigen space from **M** matrix. So the **p** can also be projected into the same eigen space. We know that all the elements in the **M** is non negative, so based on the Perron-Frobenious theorem, there exists a dominant eigen value in M matrix, which provides the first eigenvalue *λ*_1_ is positive and largest, thus guaranteed that *λ*_1_ > |*λ*_2_| > … > |*λ*_*n*_|.

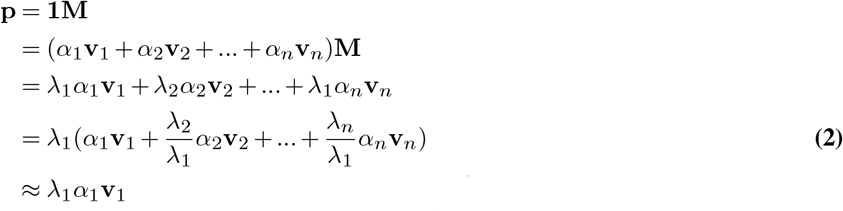

So we can also get that, the summation of **p** matrix approximately can be rewritten correlates with the first eigen value.

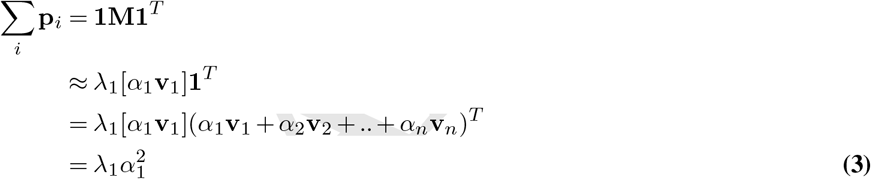

Combine with equation 2 and 3, we can see that the term removed from M matrix is actually the first eigen mode.

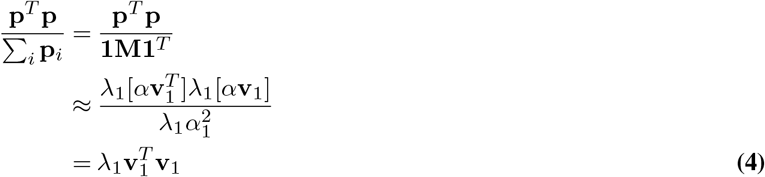

### Proof of gradient of LH equals APC

Given the definition of eigenvalue, wen can get the equations as follows.

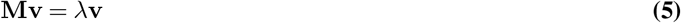

And the differentiate of the equation is

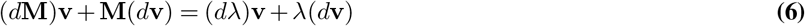

then times **v**^*T*^ at both side,

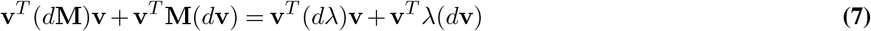

Because **M** is an hermitian matrix, you can also get **v**^*T*^ **M** = *λ***v**, and since **v** is a unit vector. So, **v**^*T*^ (*d***v**) = *d*(||**v**||^2^) = 0. Thus, we can get

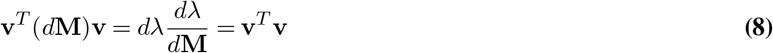

In element level, we can get

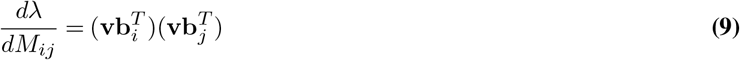

**b** is an Euclidean basis vector with a 1 at entry i and zeroes elsewhere. So the gradient of LH equals

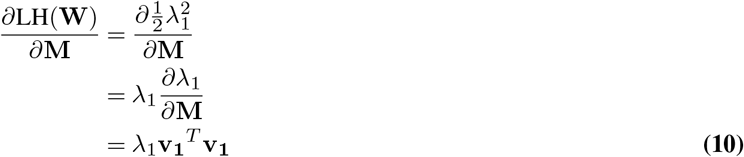

### Explanation of different regularizers

In this paragraph, we will present the differences in the eigen-space. For L2 regularizer, it is re-scaling the **M** matrix, which also means it scales all the eigen-modes.

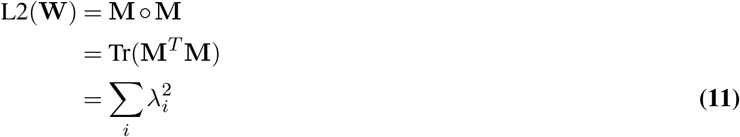

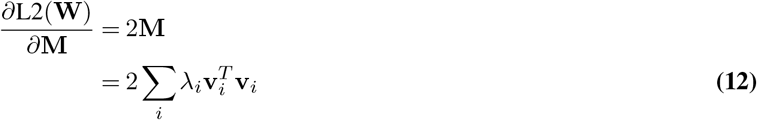

For Block L1 regularizer, it applies same gradient to each element. in eigen space, the gradient for each mode is dependent with linear coefficient sets to represent **1** vector by eigen-vectors. Thus makes it hard to tune the hyperparamter for each proteins.

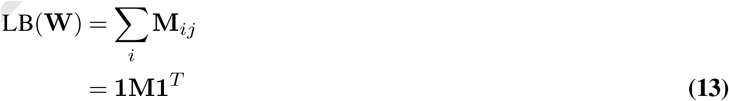

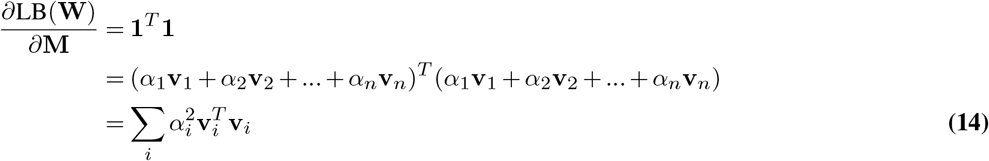

While for LH, it is already showed in equation 10, that the gradient equals to the first mode, that keeps the rest eigenmodes unchanged which can well represent the sparse signals.

## Additional Analysis

In the case of chorismate mutases, the experimentally tested fitness (denoted as “norm r.e.”, which does not necessary mean stability) of sampled sequences first showed poor correlation with the bmDCA statistical energies, but then rescued by a binary logistic regression model based on experimentally labeled natural MSA(20). The performance is also poor in L2/LH model when following the same procedure. We suspect the natural MSA contains paralogs of alternative functionality that “pollute” the bmDCA model, and guide bmDCA sampling to generate sequences of alternative or no function. So we filtered the training MSA, define norm r.e. < 0.25 as negative sequences and the rest as positive sequences, then we retrained L2, LH and bmDCA model with only feeding positive natural sequences. After that, we revalued the statistical energies of reported sequences with new trained model. Now as we can see in figure S4, the models can well separate the sequences in all of the three models. For our understanding, fitness is a high level metric which needs a model to incorporate at least the structure and function information, which might be more complicated to be investigated later.

---

**Fig. S1.**
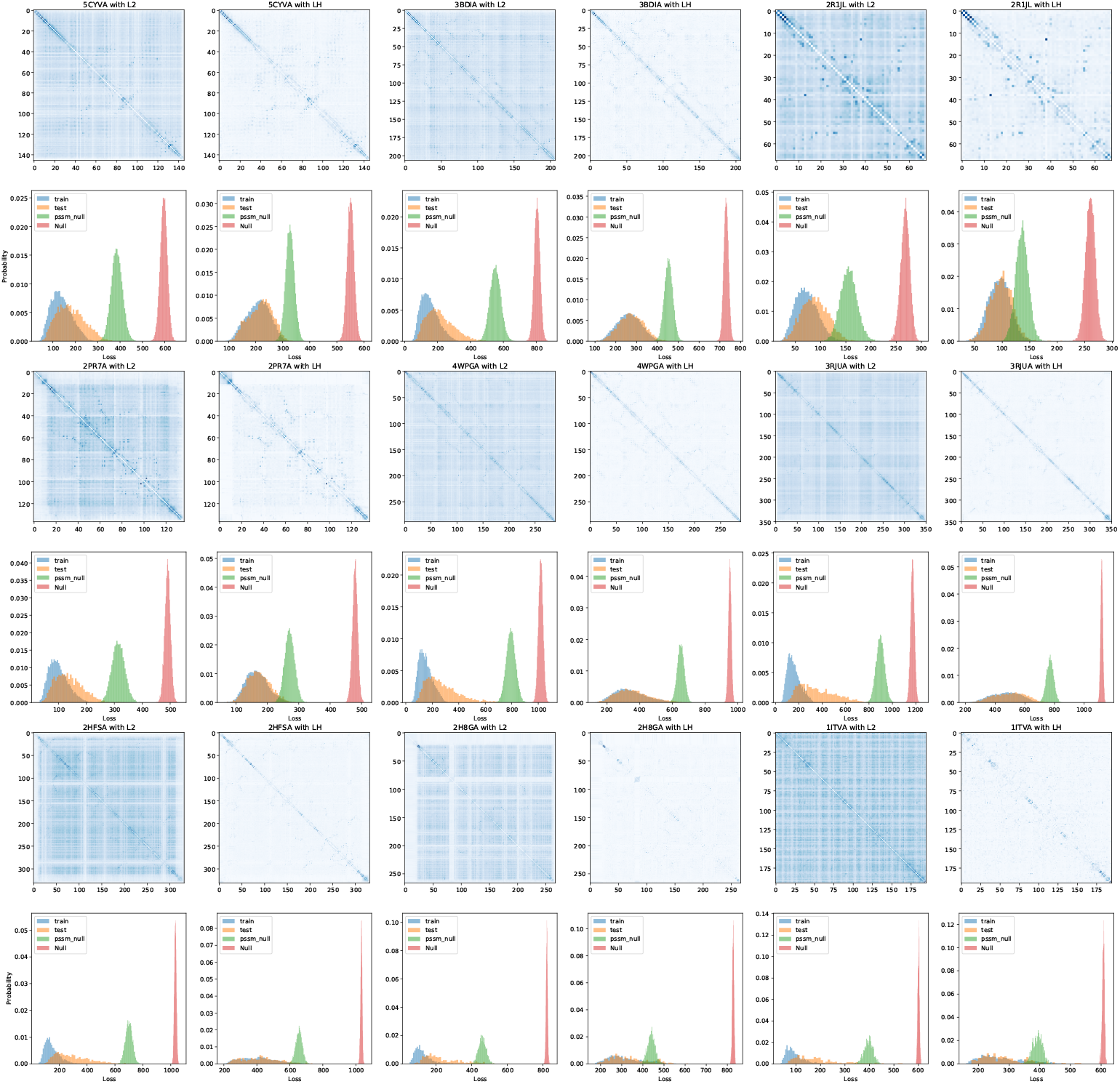
9 more proteins with various MSA depth in a 3*3 grid. In each grid, left panels shows the contact map and Loss distribution from L2 method, and right panel is from the LH method.

**Fig. S2.**
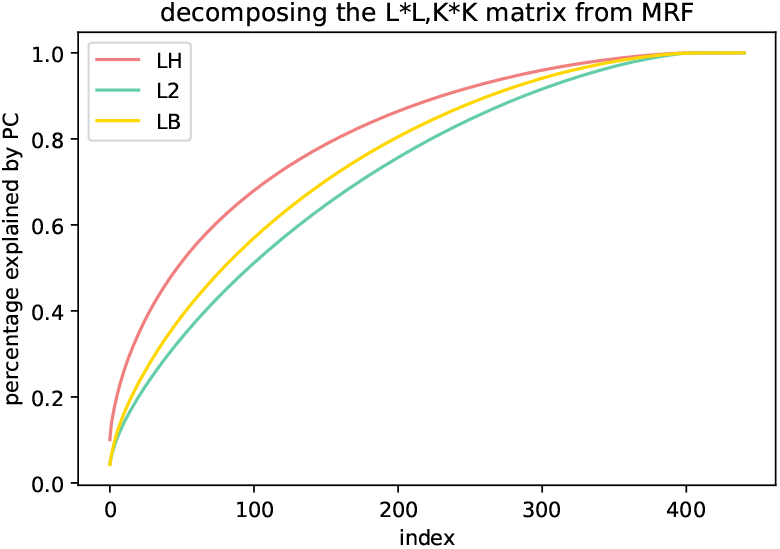
The Principle Components Analysis is applied on a L*L,K*K matrix generated from MRF, the X-axis shows index of K*K dimensions, the y-axis shows the percentage explained by principle components.

**Fig. S3.**
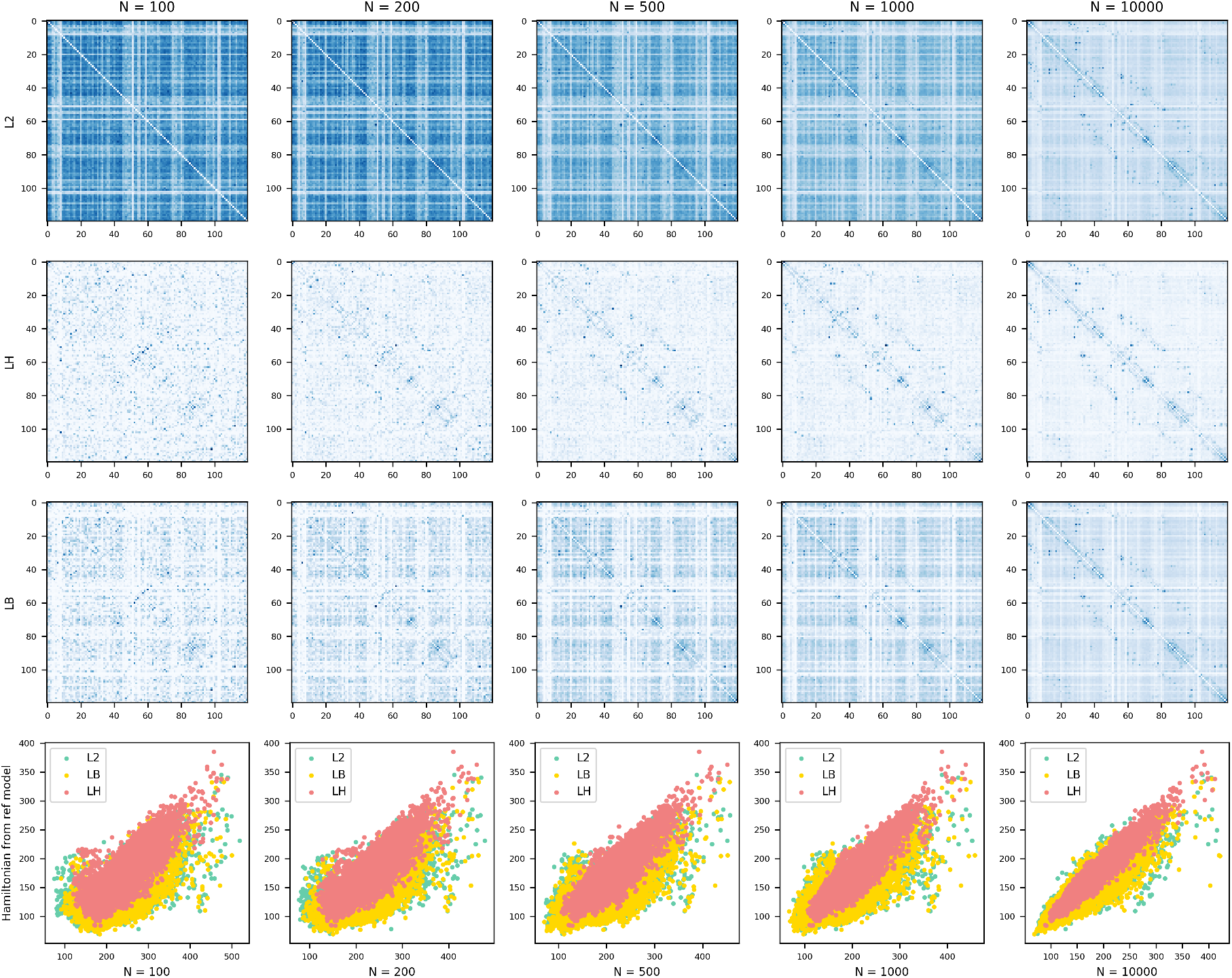
A set of experiments with various MSA depth in same protein with different regularizers. The first row is the L2 based contact matrix, second row is LH based contact matrix and the third one is LB based method, the last row is the hamiltinoian coorelation between given depth of MSA and hamiltonian infered from sufficent MSA. Green, yellow and red are used to label L2, LB and LH.

**Fig. S4.**
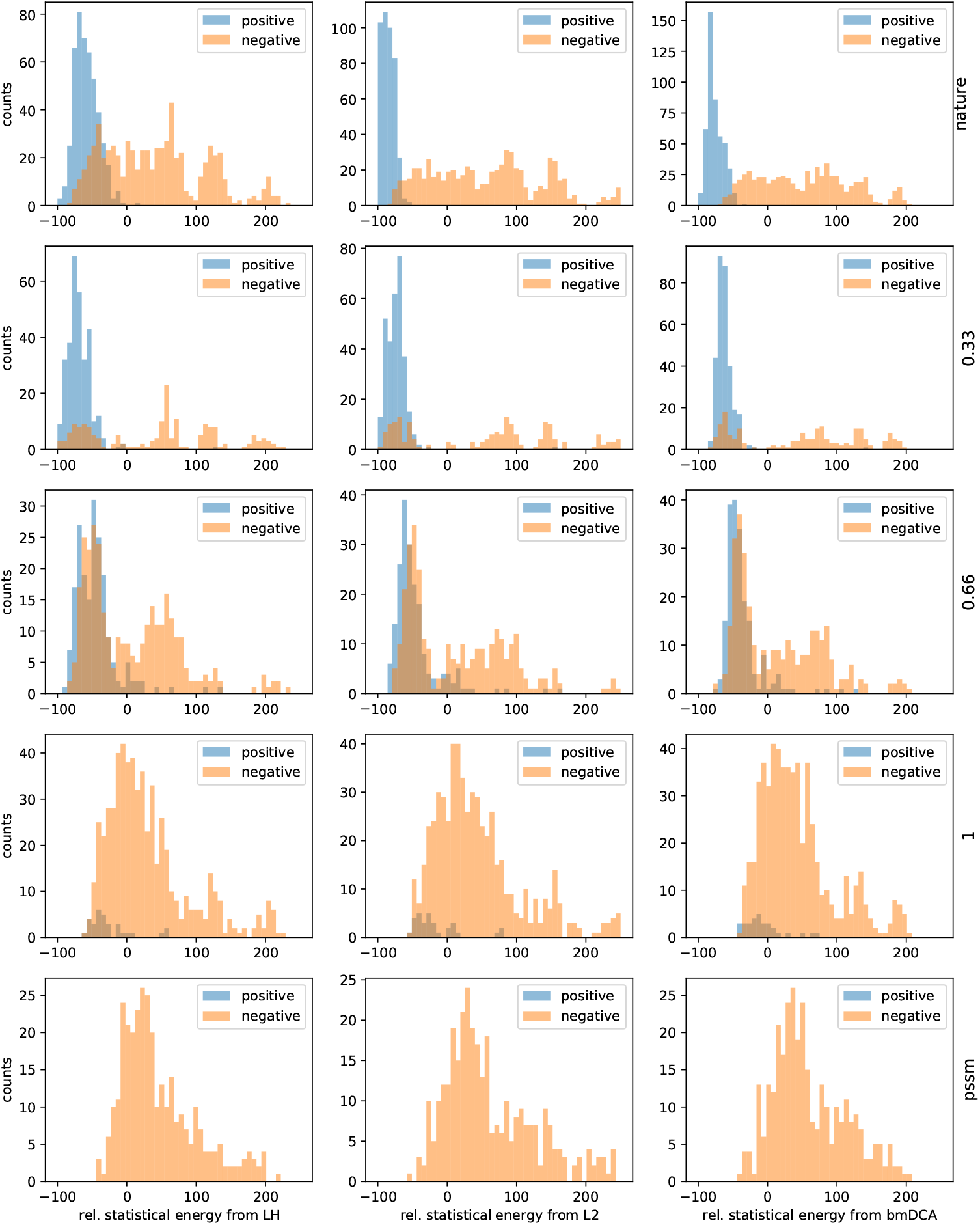
Distribution of three statistical energies for 1130 tested natural AroQ homologs reported in (20), blue and yellow labels are applied to classify positive and negative sequences based on the norm r.e. Each column shows a different regularizer. All three methods shows similar classifition ability in different sets includes nature msa and sequence generated in MC temperature with 0.33, 0.66 and 1.

**Fig. S5.**
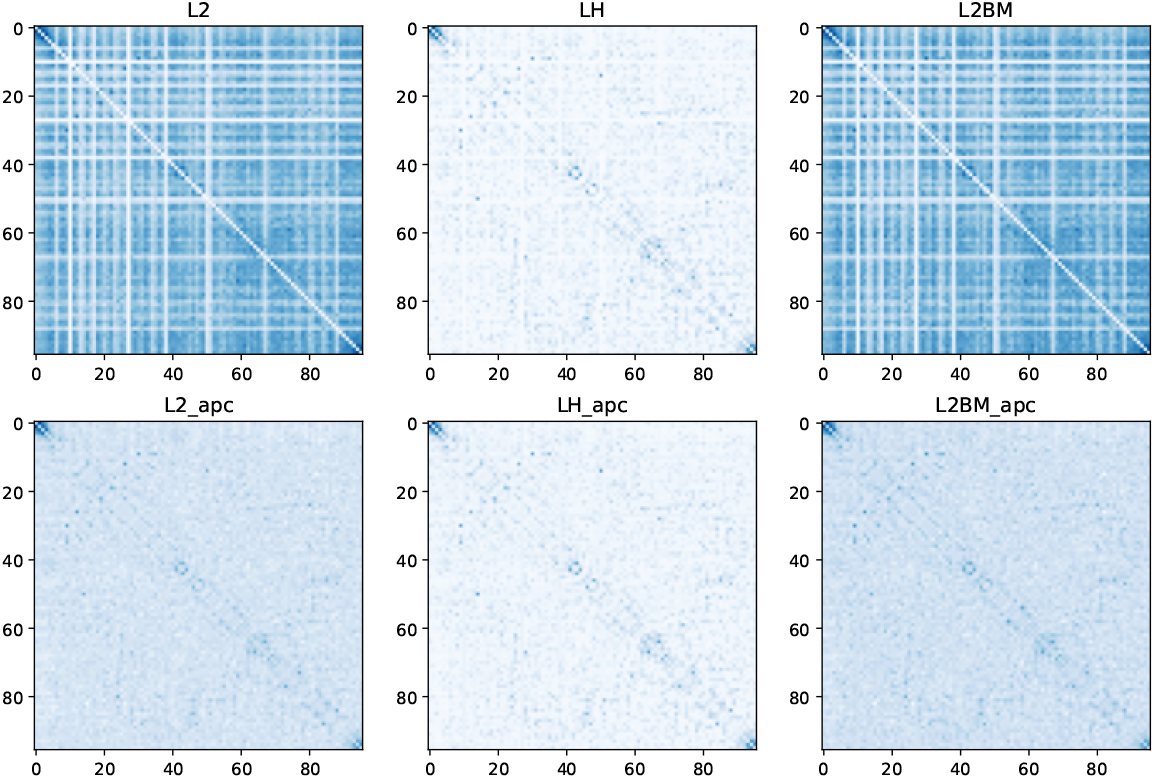
Calculating the mutual information with generated sequences by different three methods with L2 LH and BMDCA(L2BM), first row contains three co-evolutionary matrices**M** and second row is the matrices with apc correction.

**Fig. S6.**
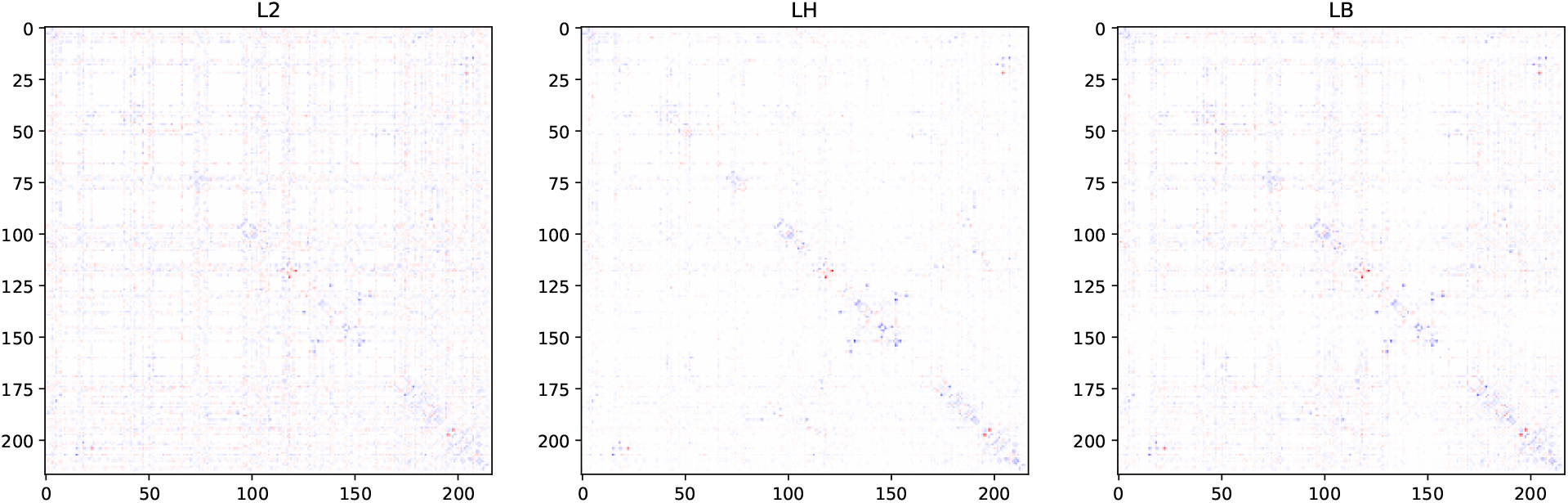
coupling strength difference between two proteins ANC1 and B.marinas. A fuzzy stripped noise can be observed from L2 and LB.

